# How minor sequence changes enable mechanistic diversity in MFS transporters? An atomic-level rationale for symport emergence in NarU

**DOI:** 10.1101/2025.11.20.689225

**Authors:** Tanner J. Dean, Jiangyan Feng, Diwakar Shukla

## Abstract

Closely related membrane transporters can diverge sharply in their modes of transport despite minimal sequence differences, underscoring how minor structural features can alter transport function. This divergence is exemplified in nitrate and nitrite transport across bacterial membranes, which supports anaerobic respiration and involves the major facilitator superfamily (MFS) transporters NarK and NarU. NarK operates as a nitrate/nitrite antiporter, whereas NarU’s mechanism remains unresolved, with evidence suggesting potential symport activity. Using extensive adaptive molecular dynamics simulations and Markov State Modeling, we mapped NarU’s conformational free-energy landscape and assessed how its behavior contrasts with mechanistic principles established for NarK. NarU follows a similar gating pathway but displays pronounced asymmetry favoring the outward-facing state and stabilizes an *apo*-occluded intermediate inaccessible to antiporters. This state arises from rotation of an arginine gating pair and a hinge glycine substitution that enhances gate flexibility. These sequence-dependent adaptations alter gating energetics and reprogram the scaffold to permit coupled co-transport. Our results show that a presence of a few strategic residue substitutions in the binding pocket and translocation pathway could alter the transport mechanism of transporters with high sequence and structural similarity.

## Introduction

Nitrate metabolism in bacteria is highly important for cell survival as a terminal electron acceptor for anaerobic respiration and has even been found involved in biofilm synthesis in Uropathogenic *E. coli* .^1^ The cycle of nitrate and nitrite transport between the cytoplasm and periplasm in *E. coli* is highly complex, with several transporters and enzymes within both the cytoplasm and periplasm known to act on these molecules. ^2–4^ Previous work has highlighted an interest in targeting nitrate/nitrite transporters as a means of evading antibiotic resistance in bacteria.^5^ While many of these transporters and enzymes have well-characterized functions, a clear understanding of their mechanisms is required when seeking to produce treatments which can avoid secondary effects on eukaryotic nitrate/nitrite transporters like the *NRT1* and *NRT2* gene families.^6^

Transport between the cytoplasm and periplasm is mediated by NarU, NarK, and NirC; of which NarU and Nark belong to the major facilitator superfamily. The MFS is a ubiquitous class of secondary active transporters that move ions and metabolites across membranes via a conserved alternating-access (“rocker-switch”) mechanism and contain uniporters, symporters, and antiporters.^7,8^ Secondary active transporters couple the movement of one substrate to the energetically favorable movement of another ion or molecule and typically follow one of three mechanisms: uniport, symport, or antiport.^9,10^ In uniport, a transporter facilitates the downhill movement of a single substrate without coupling. Symporters bind two substrates simultaneously and coordinate their translocation in the same direction, often relying on inward or outward gating asymmetries that stabilize co-bound intermediates. Antiporters instead exchange substrates in opposite directions and require a strict coupling regime that disfavors occluded *apo* intermediates and enforces alternating substrate binding. In the rocker-switch mechanism, the N-terminal and C-terminal lobes of the transporter rock around a central axis to open and close an inward-facing (IF) gate and outward-facing (OF) gate, allowing for tightly controlled transport as they cross through an IF, occluded (OC) and OF state. Nearly all MFS transporters have a conserved structure of 12 transmembrane helices, with a conserved C2 pseudo-symmetry inherent to MFS transporters. From this structure, helices 1, 4, 7, and 10 form the central cavity for transport, while the remaining helices serve as structural support. The nitrate/nitrite porter (NNP) family is a subset of MFS that includes the *Escherichia coli* proteins NarK and NarU, which facilitate nitrate and nitrite exchange to support both respiratory and assimilatory pathways. ^11^ From previous experimental analysis, NarK has been shown to operate as a nitrate/nitrite antiporter, consistent with its role in exchanging intracellular nitrite for extracellular nitrate. ^12^ Prior computational studies of NarK from Feng et al. have further extended our mechanistic understanding of this antiporter mechanism by highlighting the key residues involved in substrate recognition and exchange, as well as confirming the lack of an *apo*-occluded state as expected of a coupled antiport mechanism. ^13^

NarU is a close homolog of NarK sharing ∼75% sequence identity implying a similar functional role.^14^ Despite their similarity, NarK and NarU may not share the same mechanism. In fact, structural and biochemical studies have produced conflicting interpretations of NarU’s function ranging from strict antiporter, strict symporter, and antiporter with nitrite uptake possible under certain conditions.^15–17^ Whereas NarK’s crystal structures and liposome assays demonstrate a strict antiport cycle, the first NarU structure was interpreted by Yan et al. (2013) as a potential nitrate/(alkali cation) symporter. Due to the difficulty of long-lasting radiolabeled nitrogen isotopes,^18^ liposome assays using different ions were used to show that NarU can mediate net nitrate uptake or nitrite uptake, hinting that NarU could support symport under some conditions. This debate over NarU’s mechanism highlights a key gap in our understanding of NNP transporters and motivates our further investigation. To resolve this issue, we used extensive molecular dynamics (MD) simulations combined with adaptive sampling and Markov state modeling to map the conformational free-energy landscape of NarU and the coupling between sodium and substrate movements. Transporters regularly function on the *µ*s to s timescale and as such there are many methods that have been developed to expedite these rare sampling events^19^ such as umbrella sampling,^20^ metadyamics,^21^ and markov state models.^22^ Unlike metadynamics and umbrella sampling, markov state models do not add a bias to the simulations which must be removed and instead leverage many short simulations to build a statistical model of system dynamics. ^23–25^ MSMs have been used to investigate the transport mechanisms for different classes of membrane transporters.^13,26–31^ We built atomistic models of NarU in a complex membrane with nitrate or nitrite and explicit Na^+^ to match previous experiments^16^ and ran over 900 *µ*s of simulations using least counts adaptive sampling. By analyzing the transporter gating landscape, key substrate-protein contacts along the transport cycle, ion-substrate transport correlations, and key differences between NarU and Nark, we aimed to characterize the gating energetics and structural rearrangements that drive NarU’s transport cycle and resolve whether symport is possible.

Our simulations reveal that NarU’s gating energetics are highly asymmetric as compared to an antiporter. The IF to OC transition has a fairly low barrier (<0.5 kcal/mol), whereas the OF to OC transition is more energetically costly (∼1.6 kcal/mol). This L-shaped free-energy landscape favors the OF state and suggests that NarU readily samples the occluded state from the cytoplasmic side, an arrangement more common of symporters rather than strict antiporters.^13^ Consistent with this result, sodium and the substrate move in a concerted fashion through the pore rather than diffusing independently, indicating coupled co-transport of substrate and a driving ion. Structurally, we identify two key adaptations that promote a symport mechanism. A helix-stabilizing alanine in NarK becomes a helix-destabilizing glycine residue at a cytoplasmic helix hinge (G150) in NarU that greatly increases flexibility for intracellular gate opening and breaks the symmetry of gating energy. Additionally, a pair of conserved arginines in the binding site can adopt either a tight, substrate-coordinating configuration or a separated, *apo* conformation upon sidechain rotation to separate positive charges allowing *apo*-OC. Together, these features relax the tight gating constraints seen in NarK and allow NarU to accommodate simultaneous Na^+^ and substrate flux, as well as enter a stable *apo*-OC configuraiton. In short, our results demonstrate that subtle sequence differences such as a flexible hinge and the re-orientaiton of arginine gating residues repurpose the shared NNP fold from strict antiport to allowed symport.

## Results and Discussion

### Asymmetric gating landscapes support a symport mechanism for NarU

To evaluate whether NarU contains the L-shaped intracellular vs. extracellular gating landscape ubiquitous with rocker-switch transporters, we mapped the MSM-reweighted landscape. Figure 1A and 1B show this L-shaped landscape with no hourglass state, a state when both the intracellular and extracellular gates are open simultaneously. Additionally, the inward-facing (IF) to occluded (OC) transition for NarU involves a free energy barrier of 0.2–0.4 kcal/mol, whereas the outward-facing (OF) basin represents the deepest minimum and the OF to OC barrier is appreciably larger (1.6 to 2.0 kcal/mol) (Free energy errors are shown in Figure S1). These values indicate that, on the free-energy surface sampled in our simulations, the IF state converts readily to OC while the OF state is stabilized. Such an energetic ordering is consistent with the canonical alternating-access picture for major facilitator superfamily (MFS) proteins, in which OF, OC and IF are the principal substates and small shifts in relative free energy and barrier heights control the net transport cycle. ^9,10^

**Figure 1:**
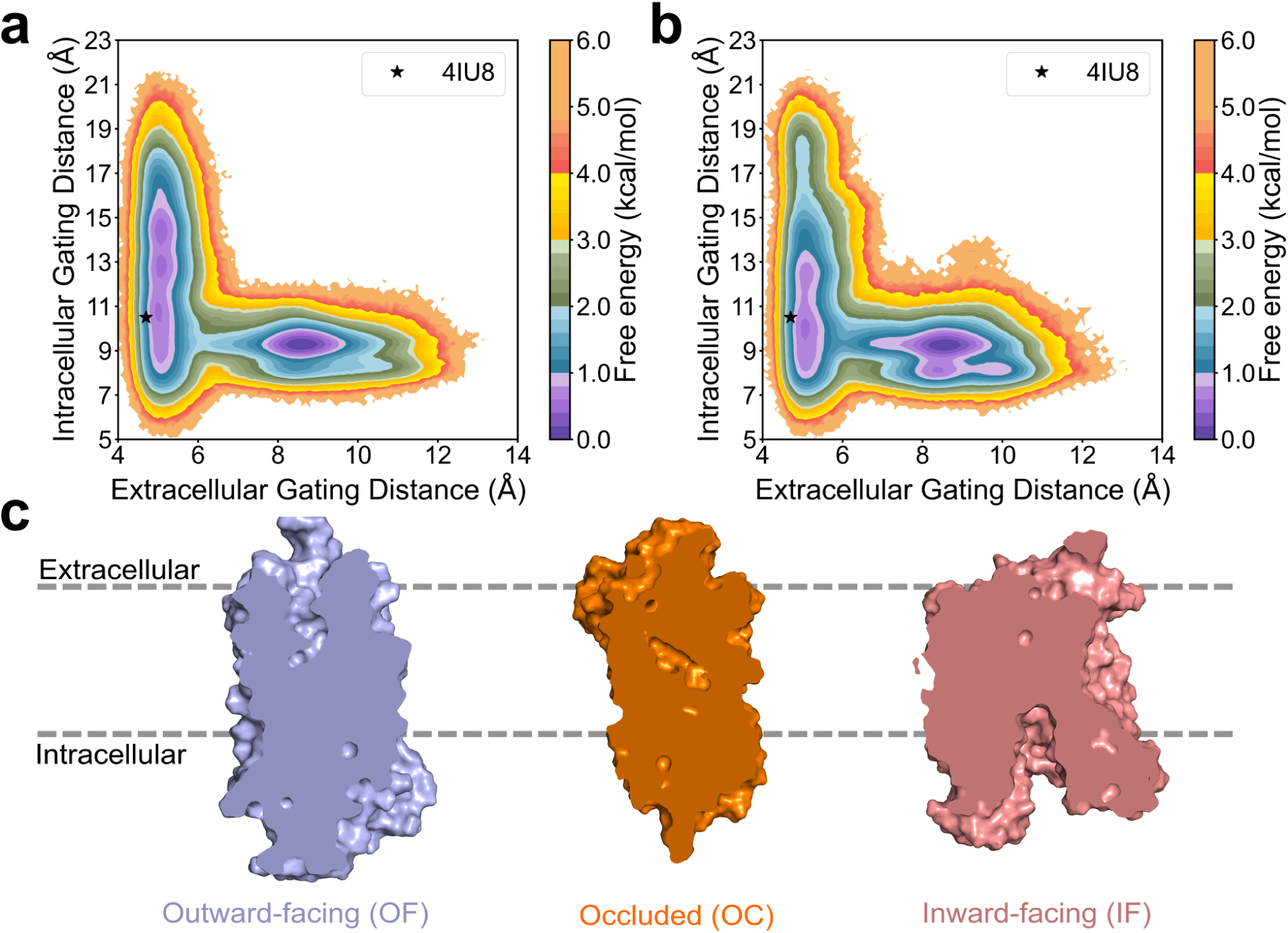
MSM-reweighted free energy landscape of the ∼900 *µ*s of simulation data was projected onto the intracellular gating distance (S44C*_α_*-A263C*_α_*) and extracellular gating distance (M139C*_β_*-F357C*_β_*) of A) NarU with NO_2_^-^/Na^+^ and B) NarU with NO_3_^-^/Na^+^ . The star represents the crystal structure of NarU chain A in PDB: 4IU8. B) Representational cross sections of the OF, OC, and IF states. The intracellular - extracellular gating distance (Δ*d*) used were 0.1 ± 0.01, 4.0 ± 0.01, and 7.0 ± 0.01 Å accordingly.

In mechanistic terms, a < 1 kcal/mol IF to OC barrier implies extensive sampling of the occluded form from the inward side in the absence of a large activation penalty; by contrast, the 1.6 - 2.0 kcal/mol OF to OC barrier corresponds to a larger energetic bottleneck. In the context of transport mechanisms, this landscape supports a symporter mechanism as these commonly rely on substrate and driving ion binding to reshape the conformational free-energy surface and bias transitions in the transport direction. Previous studies of MFS transporters have emphasized that substrate and driving-ion binding change both minima depths and transition-state stabilities,^32–34^ so an OF-stabilized *apo* state is compatible with symport behavior when substrate and driving ions are accounted for.

By contrast, recent simulation and structural work on the NarK antiporter highlights a mechanistic difference. NarK uses paired basic residues and state-dependent substrate occupancy to prevent closure of the unbound transporter and requires substrate binding to weaken electrostatic repulsion and permit conformational exchange. ^13^ In that system the IF-OC barrier can act as a rate-limiting gating step and the barrier heights between the OF to OC and OC to IF halves of the cycle are closer, producing a tightly coupled exchange mechanism with symmetric barriers. The asymmetric barriers we observe for NarU are consistent with NarU operating under a symport-like energetic regime that is less stringently substrate-gated than the NarK antiporter. In Figure 1C, a representative slice of the OF, OC, and IF states are shown demonstrating the gating entrances and occluded pocket for substrate and ion binding. The OF and IF states have been identified by extensive MD simulations.

### Localized backbone flexibility underlies sequential gate rearrangements in NarU

To map the structural elements that enable the OF to OC and OC to IF transitions, we computed residue-resolved RMSF between the OF-OC states and the OC-IF states and rendered these values onto representative backbone traces (Figure 2). The RMSF patterns display a clear spatial asymmetry between the two transitions: OF to OC dynamics are concentrated in extracellular-facing elements (notably the periplasmic ends of TM1, TM2 and are very low), whereas OC to IF dynamics shift to intracellular-facing elements (notably TM7–TM10 and the cytoplasmic loops) (Figure 2A,B). These domain-localized fluctuations indicate that NarU accomplishes alternating access through hinge-like motions of a subset of helices while preserving the central helical scaffold. Such localized rearrangements are in agreement with MFS transporters’ reliance on inter-bundle rigid-body reorientation combined with local helix bending at conserved hinge motifs.^7^ These reduced backbone fluctuation of the extraceullar loops are also consistant with prior results that re-orientaiton of sidechains at the gate and hydrophobic layer are more important for the OF transtion.^16^

**Figure 2:**
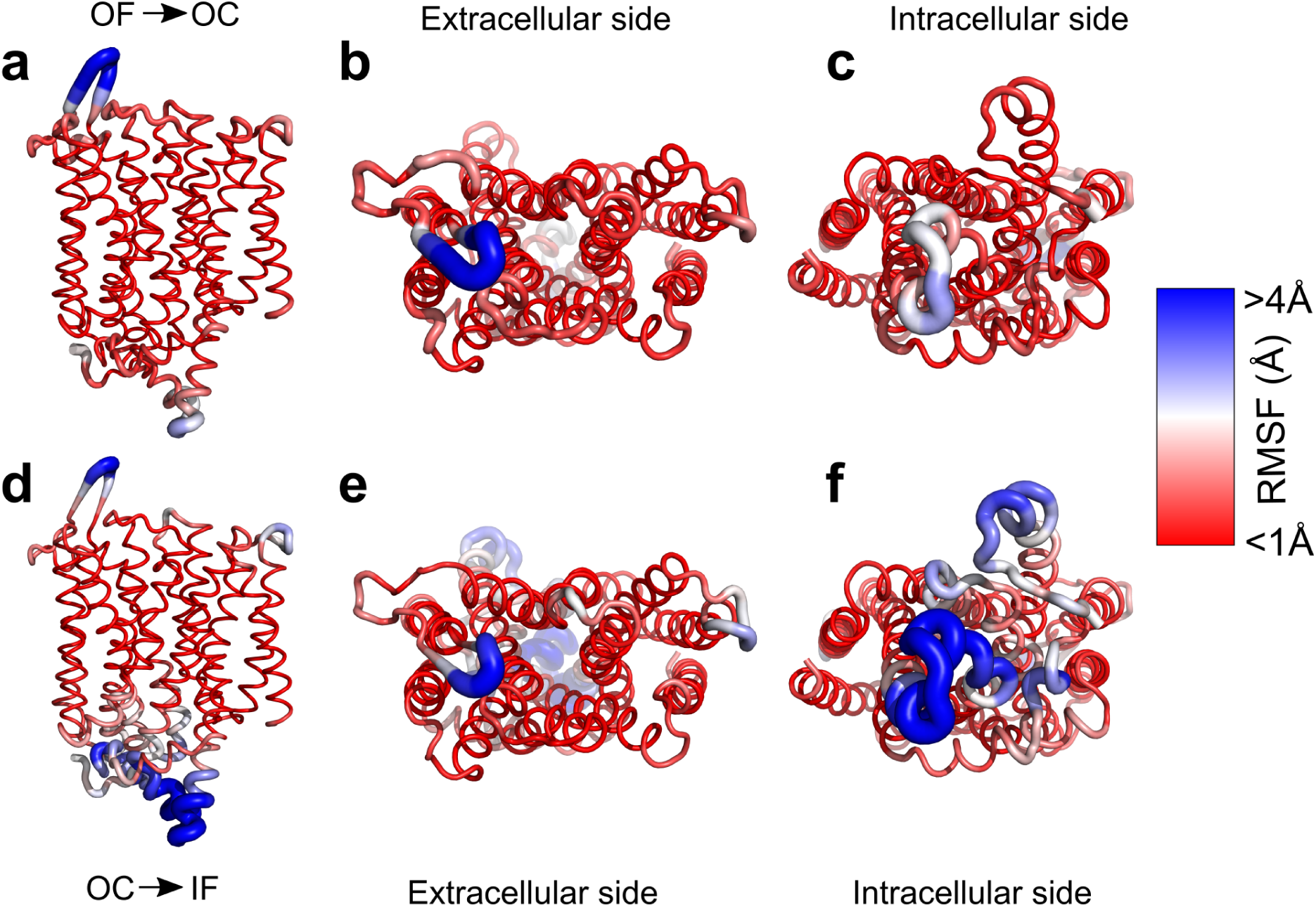
A) RMSF of the NarU backbone for the OF to OC transition from the B) ex-traceullar and C) intracellular side. D) RMSF of the NarU backbone for the OC to IF transition from the E) extraceullar and F) intracellular sides.

Two mechanistic consequences emerge from the observed pattern in RMSF of the residues of NarU. First, the dominance of periplasmic flexibility during the OF to OC transition implies that extracellular gate closure is driven by small-amplitude, high-frequency motions concentrated at the extracellular helix termini and non-backbone movements of gating sidechains; this mode is appropriate for substrate entrapment and occlusion. Second, during the OC to IF transition the amplification of cytoplasmic flexibility suggests that opening of the intracellular gate is kinetically gated by rearrangements in a distinct helix subset which reduces the probability of transient dual-open hourglass states and thereby preserves the coupling between driving ion and substrate flux. Together, these findings provide a structural basis for the sequential gating kinetics inferred from the MSM free-energy surface (Figure 1A).

At the residue and motif level, the RMSF hotspots overlap with positions that, in related transporters, participate in salt-bridge switching and hydrogen-bond rearrangements that act as molecular latches during the transport cycle. For example, the transition-specific fluctuation of the intracellular cluster (TM7-TM10) shown in Figure S2A is analogous to the cytoplasmic gating elements described in NarK and other MFS members, where coordinated disruption and re-formation of inter-bundle ionic contacts synchronizes gate motion to substrate/ion occupancy.^13^ This observed coupling between local flexibility and putative electrostatic latch elements is mechanistically consistent with the requirement that Na^+^ binding and release modulate the kinetics and thermodynamic stability of specific conformers in sodium-coupled transporters, matching experimental confirmation of sodium’s role in substrate transport.^16^

### Hydrophobic occlusion and polar contacts govern the ordered NarU transport cycle

We next sought to resolve the major gating elements for the NarU transport cycle. Figure 3 presents the position resolved free-energy projections that separate regions of high contact probability between the substrate and transporter, followed by structural snapshots annotated with residues that form the OF gate, the hydrophobic occlusion layer, and the central anion/ion coordination site for the NO_2_^-^/Na^+^ and NO_3_^-^/Na^+^ transport cycles. Free energy errors for Figure 3A and 3B can be found in Figure S5. Integrating this data yields a residue-level mechanistic map for coupling anion and Na^+^ translocation.

**Figure 3:**
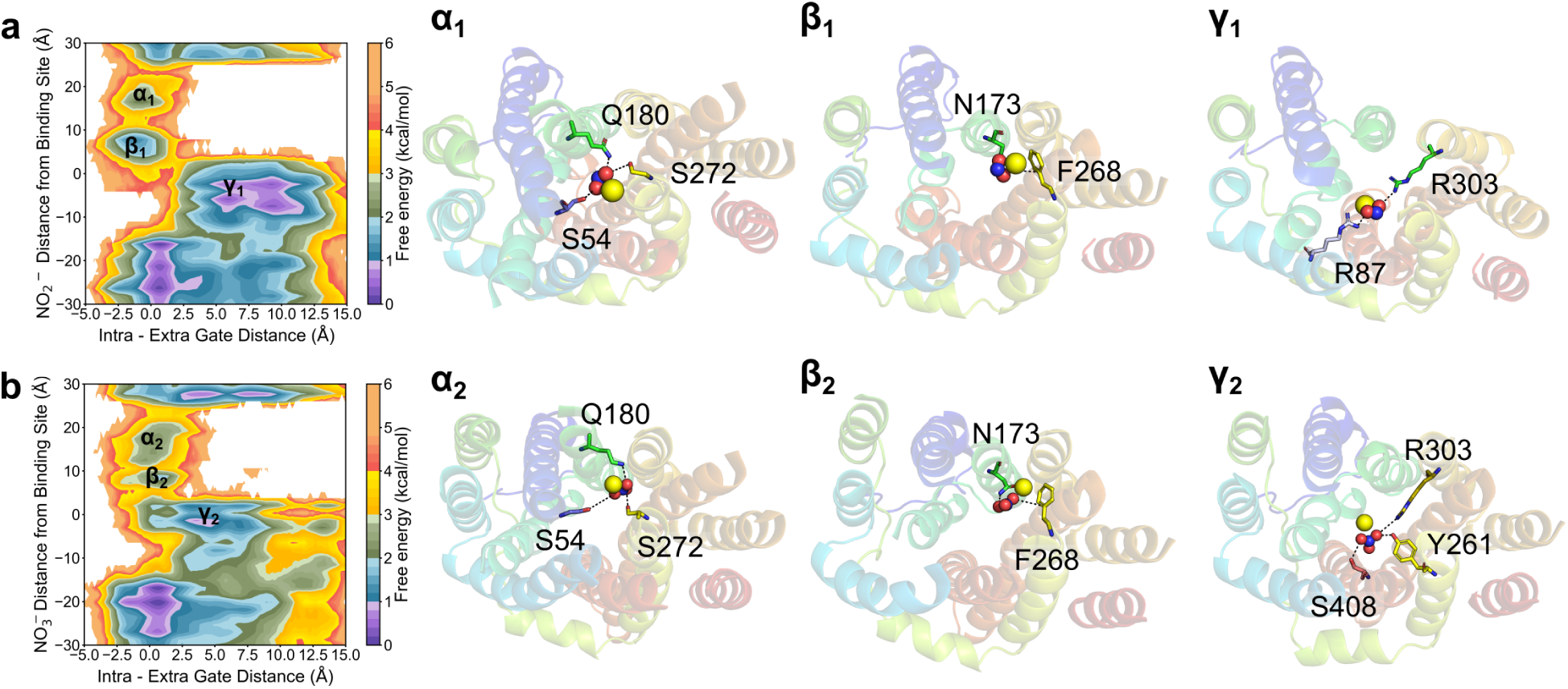
MSM-reweighted free-energy landscape of A)NO2- and B)NO3- distance from sub-strate binding site vs. Intra-Extra gating distance. α) Polar contacts with the substrate and driving ion at the OF gate. β) Contact residues at the hydrophobic gate, leading to the highest barrier between the OF and OC states. γ) The arginine gate and contacts which coordinate the substrate and driving ion at the IF gate. Quantitative contact frequencies with error are provided in Figure S3 and S4 for NO2- and NO3- respectively.

First, the NarU transporter undergoes initial anion capture and substrate binding at the extracellular gate. As shown in Figure 3*α*_1_, *α*_2_, polar side chains (e.g., Q180, S54, and S272) engage substrate oxygens within the OF basin and early OC approach. These residues form a hydrogen-bonding interface that transiently stabilizes anion entry and reduces desolvation cost as the anion is drawn from bulk solvent into the central pocket. Such polar gating residues serve as the first substrate recognition elements and are part of the extracellular gate similar to those reported for other MFS-fold transporters that guide the substrate into the binding cavity.^7^

Second, the hydrophobic occlusion layer acts as a substrate restrictive barrier. Immediately below the extracellular gate, a cluster of uncharged/hydrophobic side chains (notably N173 and F268 as highlighted in Figure 3*β*_1_,*β*_2_) forms an occlusion barrier. This uncharged/hydrophobic clamp reduces solvent penetration and creates an energetic barrier that stabilizes the occluded state by burying the ligand in a low-dielectric microenvironment. An occlusion layer is a recurring feature across diverse transporters where it functions to (a) mechanically sever extracellular and intracellular solvent pathways during occlusion, and (b) modulate the local dielectric to strengthen electrostatic interactions between the bound anion and proximal basic residues.^7,35^ The presence of this layer in NarU explains substrate preference as the substrate with driving ion must be capable of crossing this layer.

Third, binding-site polarity orchestrates the substrate coordination relay. At the central binding site (Figure 3A*γ*), charged residues (R87 and R303) coordinate the anion’s negative charge. These basic contacts provide direct electrostatic stabilization for the anionic substrate. In proximity to these anion-stabilizing residues, serine and tyrosine side chains (Y261 and S408) form a hydrogen-bond network that can coordinate or position an Na^+^ ion. This interlocking ion–substrate architecture is the canonical mechanism by which secondary transporters couple downhill ion movement to uphill substrate translocation.^7,32^ The driving ion binding remodels the local energy landscape to favor the OF (or OC) state for substrate capture and then, upon occlusion and conformational change, the coordinated release of the driving ion biases the equilibrium toward substrate release.

Overall, key contact dynamics imply an ordered symport cycle. By evaluating the key contacts underlying the OF to OC to IF cycle, we find three distinct free energy minimas along the transport path as shown in Figure 3A and B. Initially, extracellular polar contacts (Q180/S54/S272) are formed early during capture, followed by the hydrophobic/uncharged F268/N173 acting as a hydrophobic gate, and finally that central basic residues (R87/R303) engage the anion most strongly just prior to intracellular gate opening. This cycle ordering of extracellular capture followed by hydrophobic occlusion and towards ion-substrate centralization leading to intracellular release is a mechanistic sequence compatible with the symport mechanism. We next aimed to investigate whether coordinated movement of Na^+^ and substrate facilitates faster translocation compared with uncoupled steps.

### Coordinated ion-substrate binding accelerates transport

The mean first-passage time (MFPT) profiles in Figure 4 indicate that the kinetically rate-limiting step for the NO_2_^-^/Na^+^ *holo*-cycle is the *holo*-OF to *holo*-OC transition while for the NO_3_^-^/Na^+^ *holo*-cycle it is between the *holo*-OC to *holo*-IF and the *holo*-OF to *holo*-OC transitions. We further show that the direcly-coupled entry of substrate/Na^+^ is more favorable to sequential binding, with times of 19.6 and 41.8 *µ*s compared to 142 and 360 *µ*s for unpaired NO_2_^-^ and NO_3_^-^ ions respectively. This result is consistent with a transporter in which ion entry through a narrow, largely hydrophobic channel imposes a high energetic and entropic penalty that dominates the overall timescale for productive transport. The coordinated binding of the driving ion with the substrate allows for faster binding due to electrostatic pairing between the ion and substrate which partially neutralizes the net charge. This lowers the electrostatic barrier associated with crossing the hydrophobic gate, which facilitates more rapid stabilization within the binding site.

**Figure 4:**
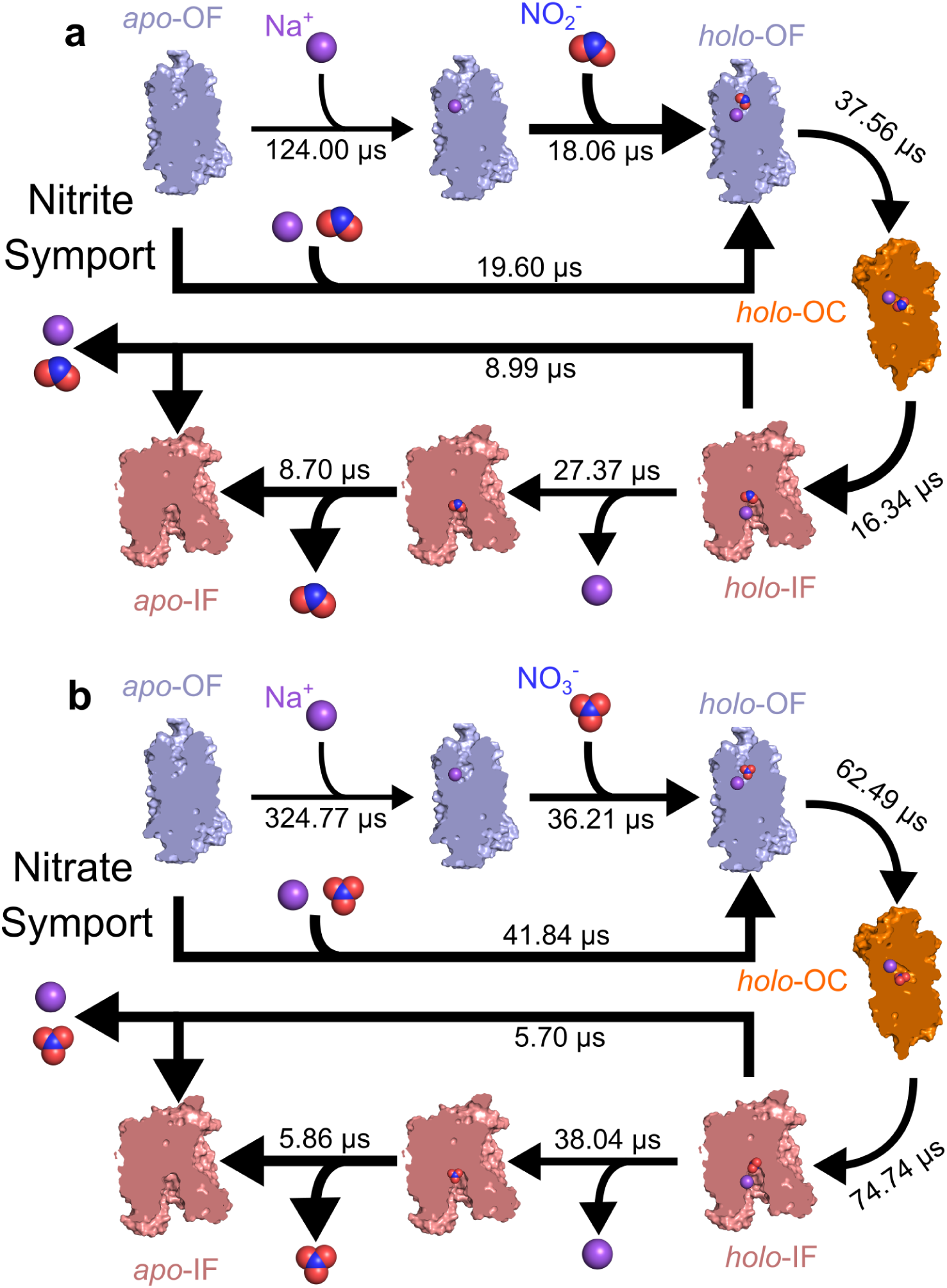
Mean first-passage time (MFPT) of the transitions between OF, OC, and IF for the A) nitrite/sodium transport system and B) nitrate/sodium transport system of NarU.

The observed slower MFPT for the NO_3_^-^/Na^+^ cycle relative to the NO_2_^-^/Na^+^ is mechanistically interpretable in terms of stronger substrate binding. Tighter substrate coordination in the binding pocket will increase residence time and thus the MFPT for completion of the *holo*-cycle. This observation aligns with independent isothermal titration calorimetry (ITC) experiments indicating higher affinity for NO_3_^-^ vs. NO_2_^-^ in NarU (33*µ*M vs. 373*µ*M).^16^ In other words, stronger substrate binding lengthens the dominant MFPT contribution arising from the bound state and the subsequent transitions required for full translocation.

The MFPT results support the existence of a symport transport mechanism biased toward co-transport of Na^+^ and a substrate as their coupling allow for passage through the hydrophobic layer: if Na^+^/substrate must bind together to open the pathway and permit substrate progression, then the transporter’s kinetics favor coupling rather than purely alternating independent exchange as expected of an antiporter. This kinetic gating explains how NarU can behave as a symporter (co-transport) rather than a strict antiporter. The occupancy and timing of the Na^+^/substrate binding event impose correlated transitions for both solutes. We further investigate this kinetic interpretation of coupled transport below.

### Correlated ion and substrate dynamics reveal symport-like transport in NarU

We next sought to determine whether this kinetic interpretation of coupled transport was consistent for both the systems (Figure 5). Figure 5A, B display the free energy landscape of NO_3_^-^ and NO_2_^-^ distance from the substrate binding site respectively vs. Na^+^ distance from the substrate binding site with free energy errors shown in Figure S6. As shown, when the substrate is under transport across the membrane (-20 to 20), there is a clear diagonal pathway demonstrating the coupled transport of the substrate with the driving ion. This co-movement of Na^+^ and NO_2_^-^/NO_3_^-^ can be quantified by computing the time-lagged cross-correlations of ion/substrate z-coordinates in bulk solution and inside the transporter. Figure 5C shows the strong positive correlations approaching 1 at *τ* =0 when the ion/substrate are within the translocation pathway compared to when in bulk solution indicate concerted motion consistent with coupled translocation. The two ions do not diffuse independently through the pore but instead display correlated positional dynamics indicative of mechanical or electrostatic coupling. This behavior is the signature of a symporter protein in which conformational changes and electrostatic networks link the motion of the substrate to the driving ion.

**Figure 5:**
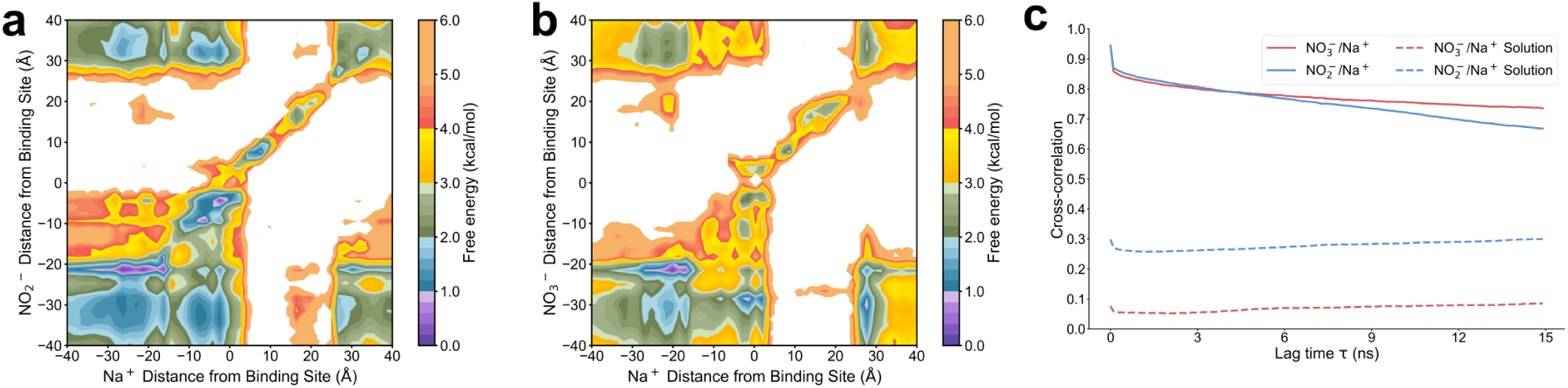
MSM-reweighted free energy landscapes of the substrate distance to the arginine gate vs. Na+ distance to the arginine gate for the A) NO2- and B) NO3- systems. C) The time-lagged cross-correlation of substrate and ion z-coordinate when crossing the membrane vs. in solution. At τ =0, the cross-correlation of z-coordinate for both systems is nearly 1 as compared to 0.2 and 0.3 in solution; implying direct coupling of transport between the substrate and driving ion.

When compared to existing structural and functional studies of NarK (a closely related NNP family member), the cross-correlation patterns in NarU differ in ways that are consistent with functional divergence. NarK has been demonstrated experimentally and structurally to operate as an exchanger (antiporter), with discrete inward-facing and outward-facing anion occupancy that favors one-for-one exchange rather than simultaneous co-transport. ^12,13^ The literature describes NarK transitions and binding site rearrangements that facilitate alternate access for single anions, whereas the correlated ion dynamics reported here for NarU implies a transport cycle in which Na^+^ binding, substrate binding, and passage are temporally overlapped rather than strictly alternating. These binding site rearrangements in an antiporter leave a O-shaped landscape in the substrate distance vs. intra-extra gating distance plot due to the lack of an *apo*-OC state. This hole does not exist in symporters, leading to an L-shaped landscape. In Figure S7, we show this separation of the O-shaped landscape of NarK antiport and the L-shaped landscape of NarU symport for both NO_2_^-^and NO_3_^-^. Thus the kinetic picture for NarU supports symport-like coupling whereas NarK exhibits classical antiport behavior.

Electrostatic coordination inside the pore by conserved positively charged residues and the ordered arrangement of side-chain dipoles can produce an energy landscape that favors co-movement. Computational and structural studies of NarK and related MFS transporters have mapped discrete anion pathways and shown how paired arginines, such as R87 and R303 in Figure 3*γ*_1_, and helix rearrangements shape barrier heights along the pathway. ^13^ Those same features can create a low-barrier corridor that carries Na^+^ together with the substrate. The cross-correlation analysis in Figure 5C provides direct evidence that the pathway in NarU supports coupled ion transit consistent via symport in contrast to the classic alternating-access antiport model established for NarK.

### Helical flexibility and arginine-gate plasticity underly symport behavior in NarU

Finally, we sought to determine what differences between NarK and NarU could potentially lead to this change in transport mechanism. Antiporters have tightly coupled opening and closing of gates aligned with alternating substrate binding for transport. Glycine at helix-loop junctions or hinge points is a recurring motif in MFS transporters that promotes backbone flexibility and lowers the barrier for helix pivoting and this has been noted in several structural and dynamical studies of MFS family members where glycine/proline residues create the necessary hinge for alternating access.^36^ While both proteins contain several conserved glycines relevant in helix bending for transport, in NarU helix stabilizing residue A150 is altered to helix-destabilizing residue G150. As shown in Figure 6A, this additional glycine allows for the breaking of the helix at the IF-gate, facilitating mobility of the adjacent IF-gate residue M149 such that the IF gate can open and close with a lowered energetic penalty. This flexibility does not exist in NarK, as shown in Figure 6B, where the lack of this G150 leads to a more rigid IF gate. A flexible G150 residue enables lower energy OC to IF and IF to OC transitions without imposing a large energetic asymmetry between states, thereby permitting conformational transitions compatible with co-transport rather than enforcing a strict one-for-one exchange cycle.

**Figure 6:**
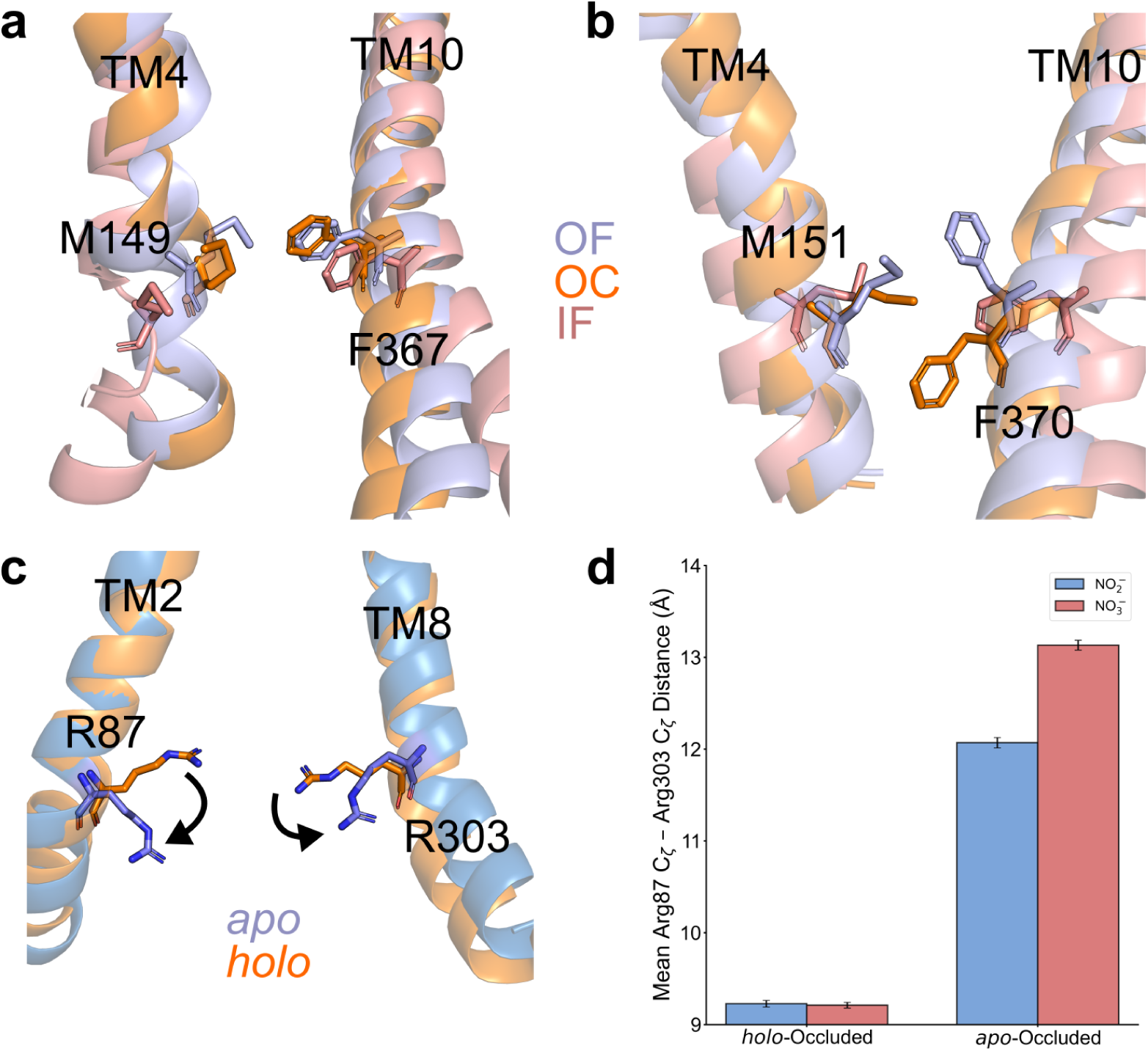
A) An A150 to G150 residue change from NarK to NarU allows for opening of the IF gate by breaking the TM4 helix. B) A150 in NarK retains a rigid IF gate as compared to NarU. C) The argine gate in NarU allows for the apo-OC state, traditionally forbidden in antiporters, by sidechain rotations of R87 and R303. D) When quantified from the holo-OC and apo-OC basins, there is a 3-4 Å change in the Cζ distance between the arginine gate corresponding with the sidechain rotation.

Additionally, it is well known that in antiporters the *apo*-OC state is forbidden, yet in Figure 3A and 3B we can clearly see the existance of the *apo*-OC state when the transporter is not directly facilitating transport. To understand why this state is allowed in NarU, Figure 6B demonstrates the distinct behavior of the binding-gate arginines that form the anion coordination site in *holo*-OC and *apo*-OC conformations. In the *holo*-OC state these arginines rigidly coordinate the anionic substrate, stabilizing the occluded substrate-bound intermediate. As quantified in Figure 6D, across both systems the zeta carbon distance between R87 and R303 was used to determine the sidechain rotation of the residues, which changes between 9.2 Å and 12-13 Å. This sidechain rotation is displayed in Figure 6C, where the sidechains of R87 on the TM2 helix and R303 on the TM8 helix lead to separation of the positive charges, and thus allow for the *apo*-OC state to exist. This conformational plasticity between a tight coordinating arrangement when a substrate is present and a relaxed, charge-separated arrangement creates permissive apo-states that are not typically allowed in strict antiporters, and thus provides further evidence for a symport mechanism in NarU.

Taken together, the G150-mediated hinge flexibility and the arginine gates ability to adopt charge-separated apo conformations offer a concrete molecular rationale for why NarU can follow a symport cycle. In antiporters such as NarK the combination of binding-site geometry and interdomain coupling enforces stricter alternating occupancy (empty occluded states are less accessible), whereas in NarU the reduced energetic gap between IF and OC and the arginine plasticity produce a landscape in which Na^+^ and substrate binding events can be temporally and spatially overlapped, enabling co-transport. These minor changes sequence and structural flexibility enable a symport mechanism separate from the known antiport mechanism of NarK, regardless of their 76% sequence similarity.

## Conclusions

This work set out to test the hypothesis that NarU, despite its high sequence similarity to the antiporter NarK, can operate through a distinct symport mechanism driven by coupled ion-substrate movement and altered gating energetics. Our molecular dynamics simulations and kinetic analyses support this hypothesis and reveal how subtle sequence-level variations translate into a fundamentally different transport cycle.

First, we resolved that NarU’s conformational energetics differ from a canonical antiporter, favoring conformations that enable simultaneous binding and co-transport. Consistent with this, the reconstructed free-energy landscape shows a stabilized outward-facing state and an asymmetric barrier profile, with a low-cost inward-facing to occluded transition and a more restricted outward transition. This landscape promotes asymmetric conformational exchange and supports coupled translocation of substrate and Na^+^, features consistent with a symport cycle.

Second, we posit that specific sequence substitutions underlie this altered gating behavior. Our analysis identifies two structural determinants that fulfill this prediction. A glycine residue at the intracellular hinge (G150) replaces a helix-stabilizing alanine in NarK, locally increasing flexibility and lowering the energetic cost of inward-gate rearrangements. In addition, the paired arginines in the binding pocket exhibit sidechain reorientation between substrate bound and *apo* states, permitting the formation of an *apo*-occluded conformation not accessible to antiporters. These adaptations collectively relax the tight gating constraints characteristic of exchange mechanisms and facilitate cooperative substrate-ion movement through the pore.

Finally, we posited that these mechanistic distinctions explain NarU’s experimentally observed capacity for nitrate or nitrite uptake under certain conditions. The simulations confirm that the combination of hinge flexibility, arginine plasticity, and asymmetric gating energetics enables a viable symport pathway consistent with these functional observations.

Overall, our findings validate the hypothesis that NarU has evolved a symport mechanism through minimal but consequential modifications to the shared NNP fold. These results define the energetic and structural features that repurpose an antiporter-like scaffold for co-transport and provide a molecular framework for understanding how even high sequence-similarity transporters can alter their transport mechanism.

## Methods

### Simulation Setup

To construct the system, we initially extracted NarU (PDB: 4IU9^37^) follwed by removing chain B (partially-IF) and keeping chain A (OC) using PyMOL (Version 3.1).^38^ To model missing missing residues 236-241 and 382-391, we used homology modeling via SWISS-MODEL.^39^ Following modeling of missing residues, the protein was protonated to match the physiological *E. coli* periplasm/cytoplasm pH of 7.5. ^40^ We next capped the termini for neutrality with the acetyl (ACE) n-termini cap and methylamide (NME) C-termini cap. NarU was next placed in a membrane bilayer generated using CHARMM-GUI.^41,42^ The membrane bilayer used a complex lipid composition of 75% PE, 20% PGR, and 5% OL/PA to best match lipidomic studies.^43^ The system was then solvated using the TIP3P water^44^ and 150 mM NaNO_2_ or 150 mM Na/NO_3_ respectively for the two systems. We avoid the addition of Cl^-^ to better match the experimental procedure we are following, as Cl^-^ is known to compete with NO_2_^-^ and NO_3_^-^ for transport.^37^ The CHARMM36m force field was used to model all atomic interactions.^45,46^ PACKMOL was used to repack the system following initial creation and minimization in CHARMM-GUI. ^47^ Hydrogen mass repartioning was used to enable the use of a 4fs timestep.^48^ The final number of atoms for the NaNO_2_ and NaNO_3_ systems were 84,469 and 84,484 with box sizes of 82 x 80 x 124 Å^3^ and 85 x 80 x 120 Å^3^ respectively.

### Pre-production MD

For all pre-production MD, AMBER22 was used.^49^ To prepare the simulations for production, energy minimization was first run using steepest descent for 5000 steps followed by conjugate gradient descent for 45000 steps. Following minimization, heating in the NVT ensemble was then conducted from 0 to 300 K for 5 ns. After heating, the system was then equilibrated under the NPT ensemble. To ensure stability, the systems were first equilibrated for 5ns at 300 K and 1 bar with all protein heavy atoms restrained via a 5 kcal/mol*Å^2^ force constant to allow the membrane and solvent to relax. Following these restrained simulations, the entire system was allowed to relax for 10 ns under no restraints at 300 K and 1 bar.

### Production Simulations

OpenMM^50^ (version 7.7) was used for production MD simulations. All production MD simulations used a timestep of 4fs and periodic boundary conditions. To maintain the temperature at 300 K, the Langevin thermostat^51^ was used with a friction coefficient of 1.41 ps^-1^. The system pressure was maintained at 1 bar using the Monte Carlo membrane barostat. The particle mesh ewald (PME) method^52^ was used to compute long-range interactions and the non-bonded interactions cutoff was set to 10 Å. Simulation frames were saved every 100 ps. Finally, all simulations were run using the Folding@Home (https://foldingathome.org) distributed computing project.^53,54^

### Adaptive Sampling and MSM Construction

Due to the long timescales associated with membrane transport events, we leveraged adaptive sampling and markov state model construction to enable more rapid exploration of potential transport events. Least-counts based adaptive sampling was performed on the inward-facing (IF) and outward-facing (OF) gates of the transporter, with all contact distances used computed by MDTraj(Version 1.9.7).^55^ While in recent years more advanced methods of adaptive sampling have been published,^56–58^ due to the lack of available anchors outside of the crystalized OC state least-counts has been previously found to be nearly always optimal for exploration. ^59^ In our least counts sampling, all frames are clustered by the IF and OF gate distances into N clusters, and the M clusters with the lowest counts are used as starting frames for the next round of simulations. While in recent years more sophisticated adaptive sampling methods have been proposed,^59,60^ for a system where we have two clearly defined gating pairs least counts performs well with low cost. In total, 921.80 *µ*s of NarU NO_2_^-^/Na^+^ and 926.60 *µ*s of NarU NO_3_^-^/Na^+^ simulations were run until the complete transport cycle was resolved. Complete sampling per round can be found in Table S1. Before construction of the MSMs, time-lagged Independent Component Analysis (tICA) was used to reduce the dimensionality of our data. ^61^ For both systems, 38 features, 36 C*_α_*-C*_α_* distances and the closest substrate and ion distance to the arginine gate, were used in tICA and MSM construction with the exact list of features found in Table S2. To optimize the parameters for our MSMs, each systems data was reduced by tICA using varied variance cutoffs and clustered by k-means clustering prior to evaluating each hyperparameter pair by their VAMP2 score using VAMPnets with the tICA plots shown in Figure S8.^62^ For both systems, the parameters with the highest VAMP2 score were used to evaluate the convergence of the implied timescales for the MSM lagtime. The best set of parameters was 900 clusters and variance cutoff 50 for Na/NO_2_ and 800 clusters and variance cutoff 50 for Na/NO_3_ in Figure S9. For both systems, 15 ns was chosen as the MSM lagtime and the full implied timescale plots can be seen in Figure S10. After construction, the MSMs were validated using the Chapman-Kolmogorov test using 5 macrostates and the deeptime python package (Version 0.4.4) in Figure S11. ^63^

### Error Analysis

Errors associated with the MSM-reweighted energy landscapes, intermediate contacts analysis, arginine-arginine gating distance, and RMSF plots were calculated by bootstrapping.^64^ For the MSM-reweighted errors, each of their variables were computed 200 times using 80% of the total data, while for the contacts analysis, arginine-arginine gating distance, and RMSF plots 10 sets of 1000 random frames from the associated basins (IF, OF, OC, etc.) were used to compute the associated values. The standard deviations of the calculated values were then used as the error. The error plots for the MSM-reweighted landscapes are located in Figure S1, S5, and S6, the contact frequences of the intermediate contacts analysis in Figure S3 and S4, and RMSF per-residue values in Figure S2.

### Trajectory Analysis and Visualization

All processing of trajectories was done using cpptraj (Version 6.4.4).^65^ All rendering of images was completed using PyMOL (Version 3.1).^38^ Substrate-residue contacts were computed using getcontacts^66^ and residue-residue distances were computed with MDTraj (Version 1.9.7).^55^ Matplotlib (Version 3.7.1) ^67^ and Numpy (Version 1.20.3) ^68^ were used for all plot development and numerical work.

## Acknowledgement

D.S. and T.J.D. acknowledge support from the National Institutes of Health Award R35GM142745.

## Author Contributions

D.S. acquired funding for the project. D.S. and J.F. conceptualized the study. T.J.D. and J.F. performed simulations. T.J.D. performed analysis. T.J.D. wrote the initial draft. D.S. and T.J.D. contributed to manuscript editing.

## Competing Interests

The authors declare no competing interests.

## Data Availability

Simulation trajectories stripped of waters and used for analysis are available upon request.

## Code Availability

Code for adaptive sampling, system preparation, and data analysis will be made available upon acceptance.

## Notes

### Competing Interest Statement

The authors have declared no competing interest.

