## Supplementary Information for "How minor sequence changes enable mechanistic diversity in MFS transporters? An atomic-level rationale for symport emergence in NarU"

Table S1: Sampling per round for both the Naru Na/$NO_2$ and NarU Na/$NO_3$ systems.

| Sampling Round | NarU Na/$NO_2$ ($\mu$s) | NarU Na/$NO_3$ ($\mu$s) |
|---|---|---|
| 1 | 99.60 | 99.70 |
| 2 | 17.30 | 20.00 |
| 3 | 2.50 | 2.50 |
| 4 | 99.80 | 99.60 |
| 5 | 20.00 | 19.90 |
| 6 | 19.90 | 20.00 |
| 7 | 20.00 | 19.80 |
| 8 | 24.90 | 25.00 |
| 9 | 19.90 | 19.80 |
| 10 | 19.80 | 20.00 |
| 11 | 19.80 | 20.00 |
| 12 | 20.00 | 19.90 |
| 13 | 19.90 | 19.80 |
| 14 | 20.00 | 20.00 |
| 15 | 19.60 | 19.90 |
| 16 | 14.80 | 14.90 |
| 17 | 19.90 | 20.00 |
| 18 | 19.90 | 19.90 |
| 19 | 19.90 | 20.00 |
| 20 | 20.00 | 20.00 |
| 21 | 49.80 | 49.80 |
| 22 | 49.90 | 49.90 |
| 23 | 50.00 | 49.90 |
| 24 | 49.70 | 49.80 |
| 25 | 19.80 | 19.90 |
| 26 | 12.50 | 12.50 |
| 27 | 12.50 | 12.50 |
| 28 | 12.50 | 12.50 |
| 29 | 19.90 | 19.80 |
| 30 | 40.00 | 39.90 |
| 31 | 39.90 | 40.00 |
| 32 | 1.40 | 1.70 |
| 33 | 4.10 | 4.10 |
| 34 | 8.20 | 9.10 |
| 35 | 9.40 | 9.60 |
| 36 | 4.70 | 4.90 |
| Total | 921.80 $\mu$s | 926.60 $\mu$s |

Table S2: Features used for tICA and MSM construction. For features 37 and 38 where there are multiple ions in solution, the ion with the closest average distance from the arginine gate for each simulation was used as the feature.

| Feature Number | Feature |
|---|---|
| 1 | T261-P367 |
| 2 | S258-I407 |
| 3 | A87-A146 |
| 4 | L262-G411 |
| 5 | A322-L361 |
| 6 | V56-L117 |
| 7 | T116-I134 |
| 8 | P53-P79 |
| 9 | L36-I167 |
| 10 | I38-M232 |
| 11 | T319-T366 |
| 12 | V40-A168 |
| 13 | T50-C138 |
| 14 | G270-I418 |
| 15 | L43-P145 |
| 16 | V104-T229 |
| 17 | C48-M179 |
| 18 | S54-A273 |
| 19 | M149-P367 |
| 20 | A87-A303 |
| 21 | A87-T261 |
| 22 | A87-A173 |
| 23 | A303-T261 |
| 24 | A303-A173 |
| 25 | A173-T261 |
| 26 | L62-L280 |
| 27 | L62-A290 |
| 28 | L280-A290 |
| 29 | P145-P367 |
| 30 | S54-G180 |
| 31 | S54-S272 |
| 32 | G180-S272 |
| 33 | P47-P265 |
| 34 | T50-P265 |
| 35 | M51-P268 |
| 36 | M51-I269 |
| 37 | Substrate dist. from arginine gate |
| 38 | Sodium dist. from arginine gate |

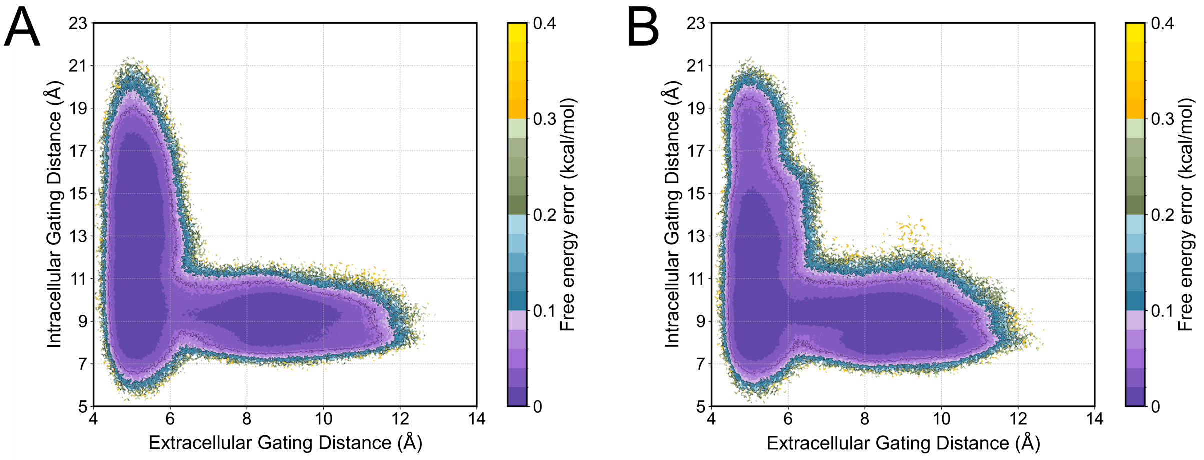


Figure S1: Error of Figure 2 gating landscape calculated by bootstrapping 80% of the total simulation data 200 times for the A) NarU $Na^+/NO_2^-$ and B) NarU $Na^+/NO_3^-$ systems.

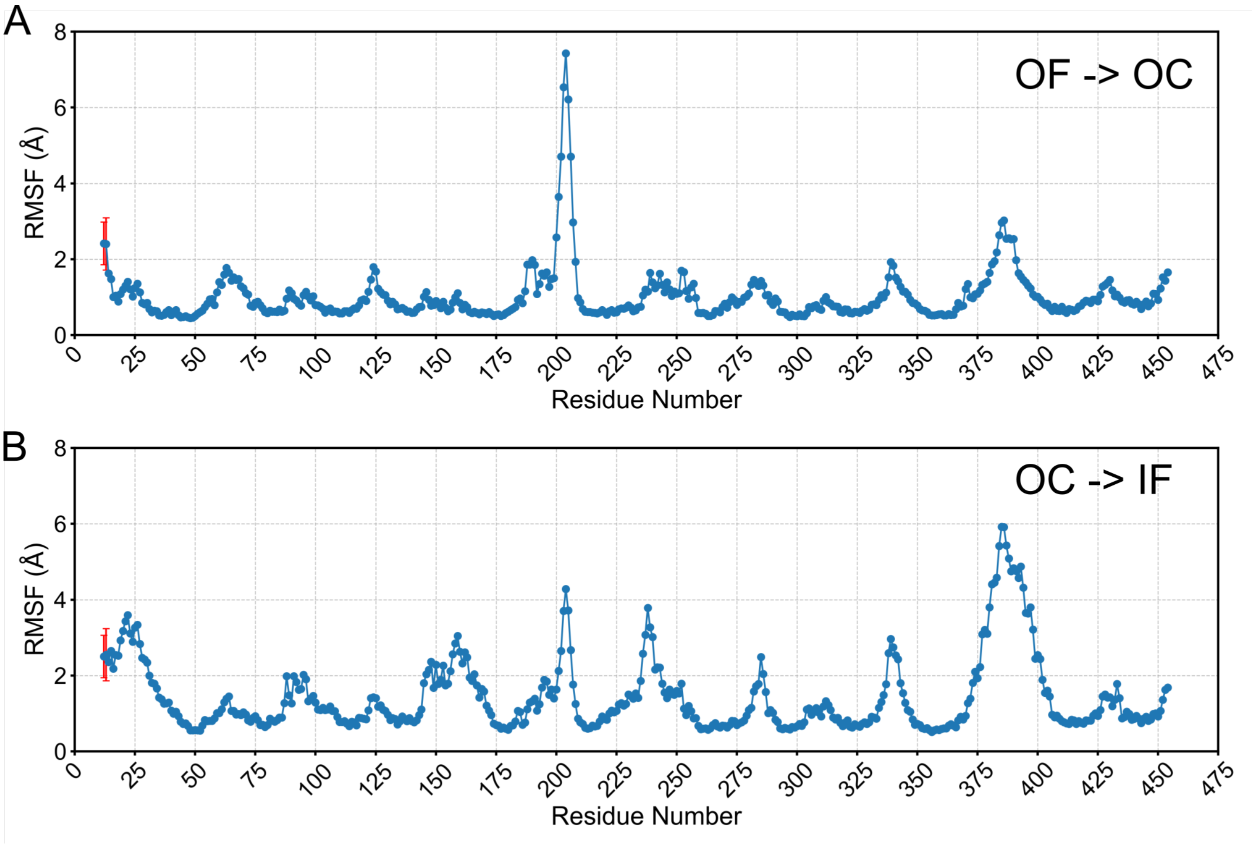


Figure S2: Per-residue backbone RMSF values with error bars for the A) OF to OC and B) OC to IF transitions of NarU. For all calculations, 1000 frames from the source were used to compute the RMSF vs. 1000 reference frames from the target. Errors are the standard deviatoins of these values.

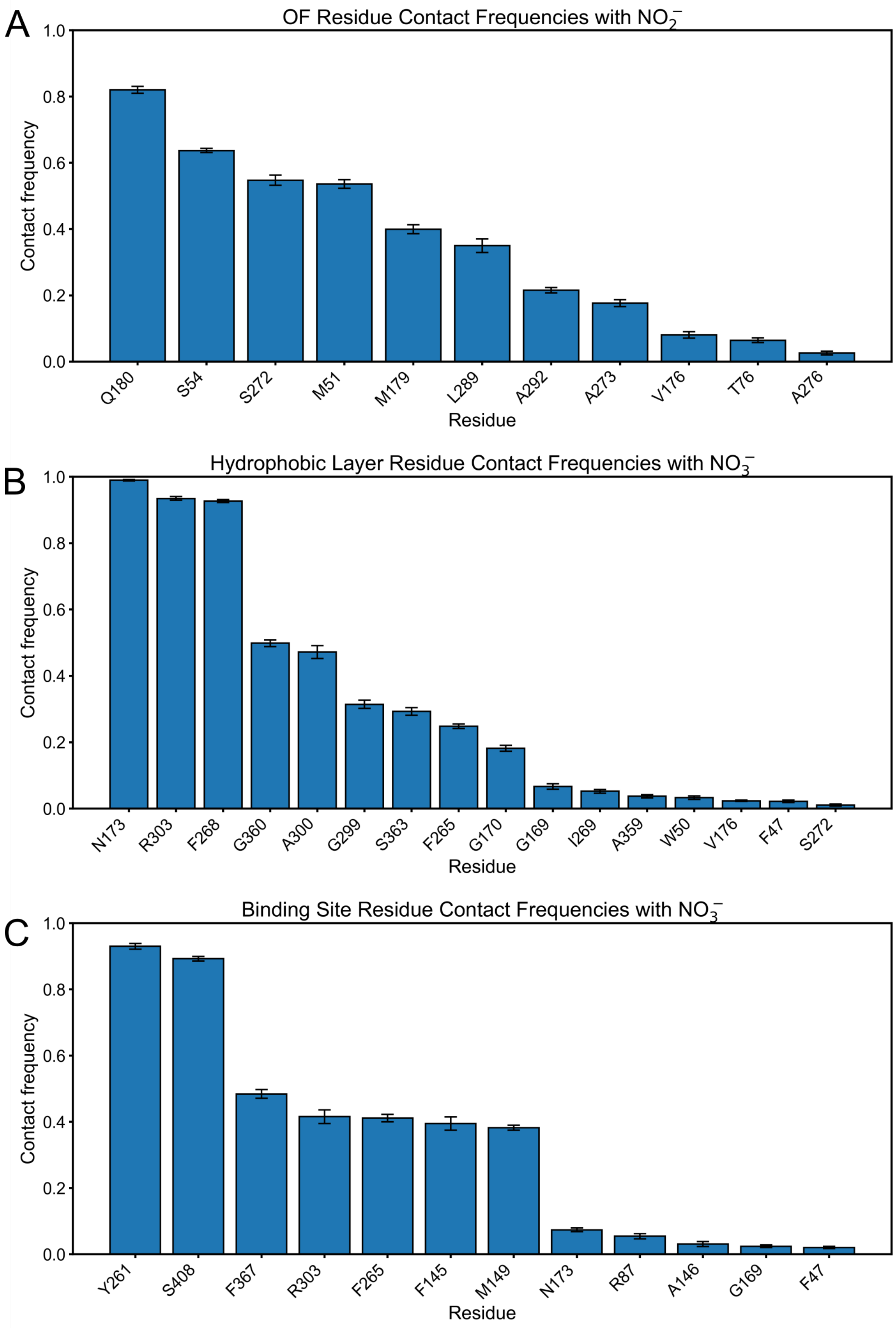


Figure S3: Contact frequecies between NarU and $NO_2^-$ computed using getcontacts for the A) OF gate, B) hydrophobic layer, and C) arginine gate. For each calculation, 10 sets of 1000 random frames from the referenced basin was used to compute the contact frequencey with the substrate. Only those residues with greater than 1% contact frequency were plotted.

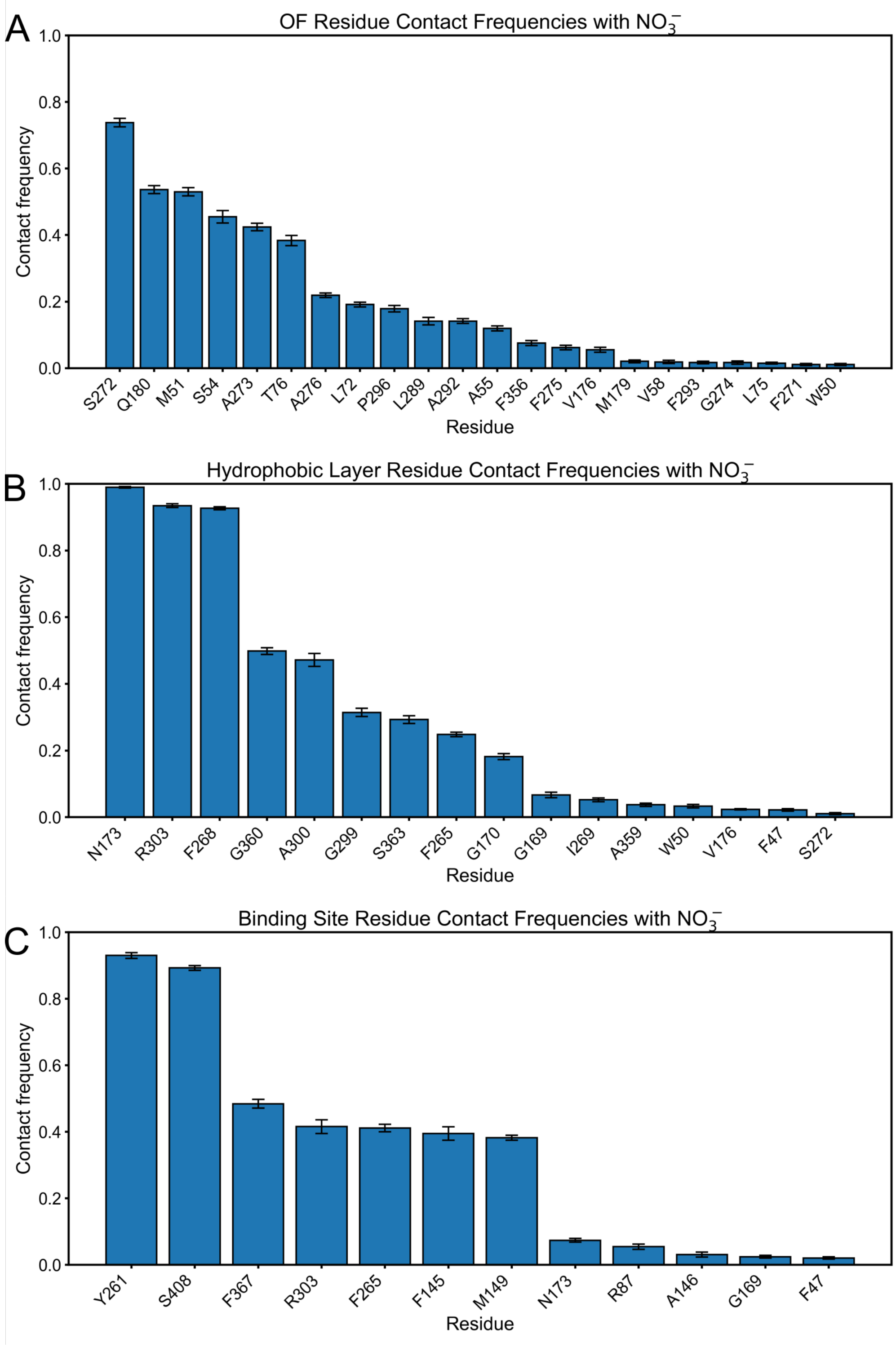


Figure S4: Contact frequecies between NarU and $NO_3^-$ computed using getcontacts for the A) OF gate, B) hydrophobic layer, and C) arginine gate. For each calculation, 10 sets of 1000 random frames from the referenced basin was used to compute the contact frequencey with the substrate. Only those residues with greater than 1% contact frequency were plotted.

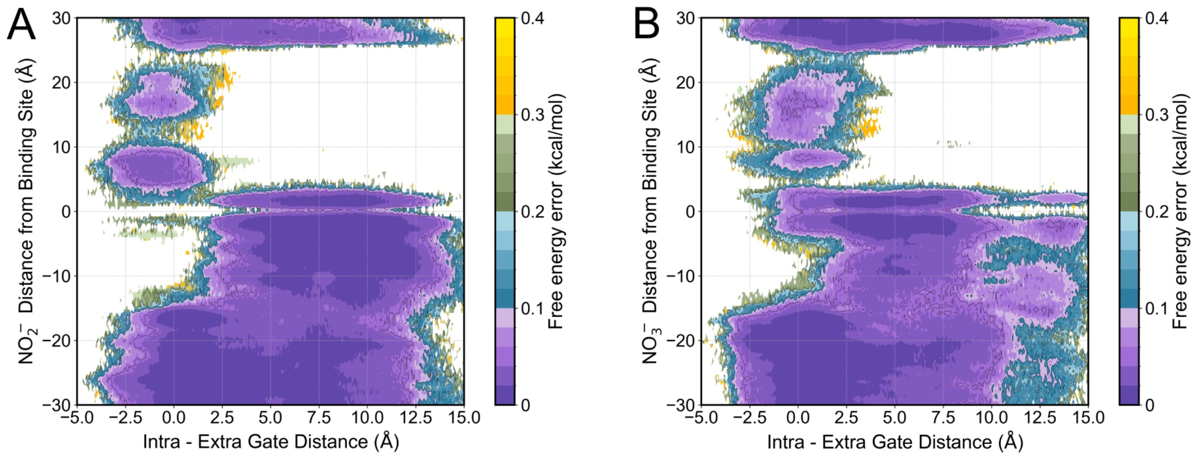


Figure S5: Error of the substrate distance from arginine gate vs. gating distance landscape calculated by bootstrapping 80% of the total simulation data 200 times for A) $NO_2^-$ and B) $NO_3^-$.

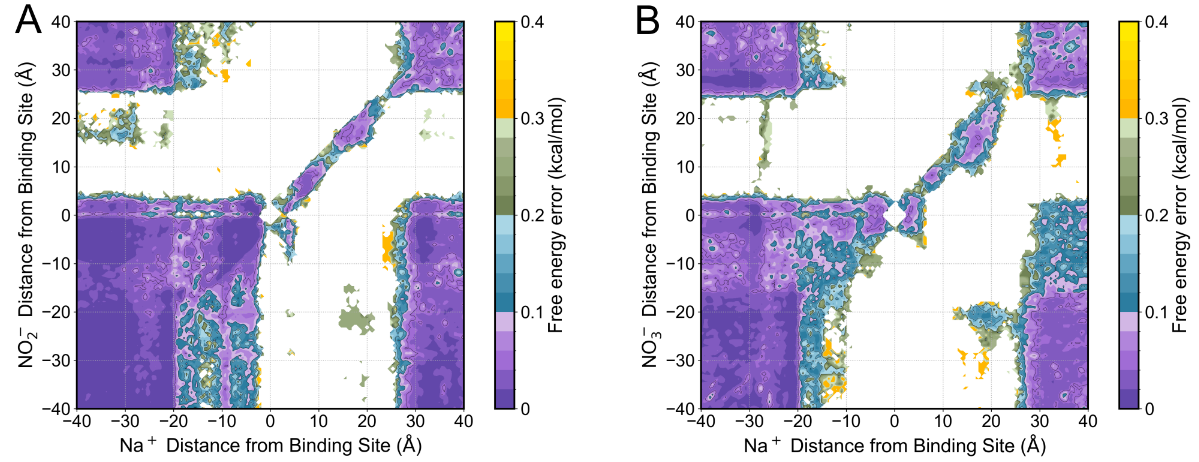


Figure S6: Error of the substrate distance from arginine gate vs. $Na^+$ distance from arginine gate calculated by bootstrapping 80% of the total simulation data 200 times for A) $NO_2^-$ and B) $NO_3^-$.

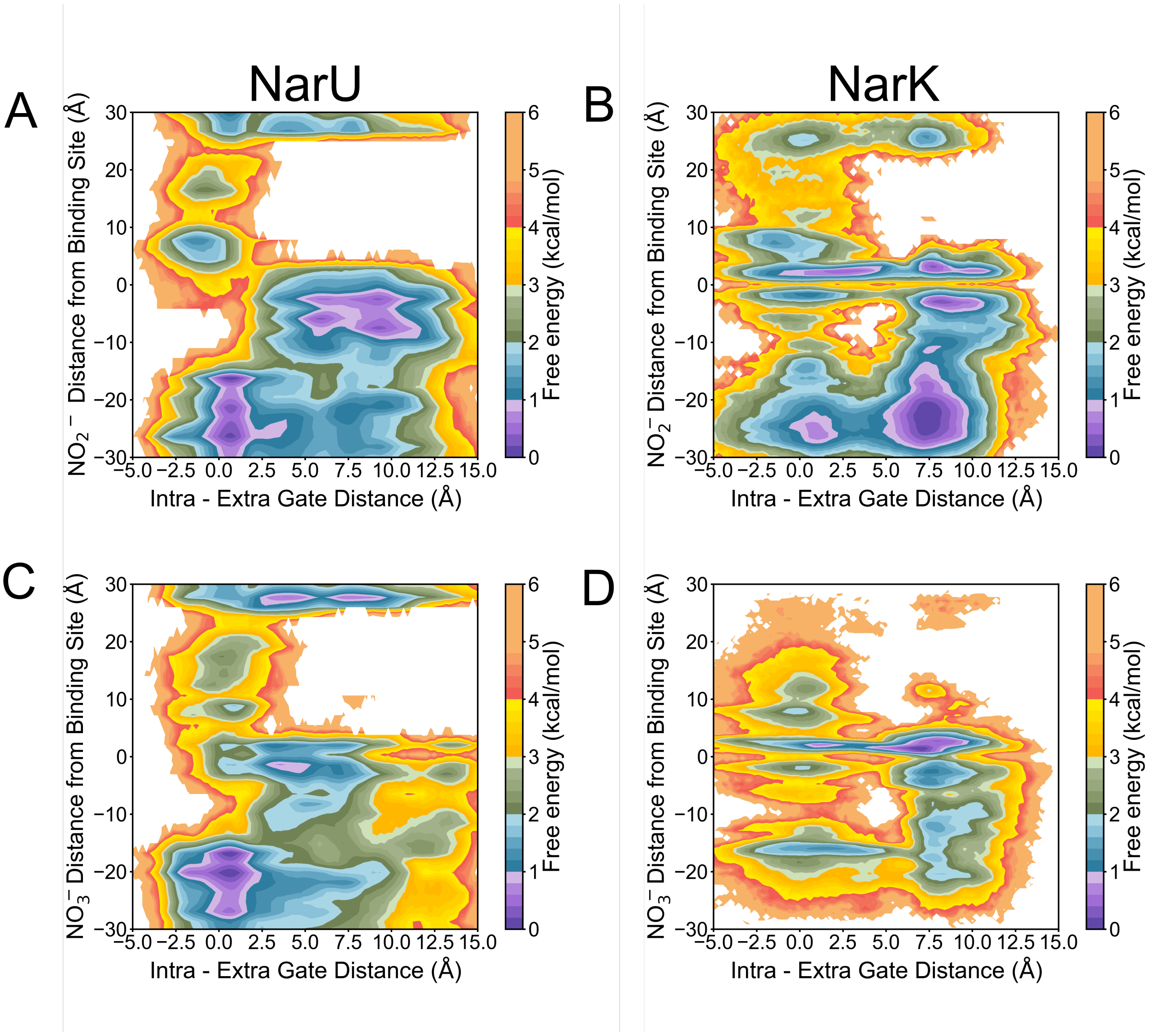


Figure S7: Substrate distance from the arginine gate vs. gating distance landscape for $Na^+/NO_2$ transport by A) NarU and B) NarK. Substrate distance from the arginine gate vs. gating distance landscape for $Na^+/NO_3$ transport by C) NarU and D) NarK.

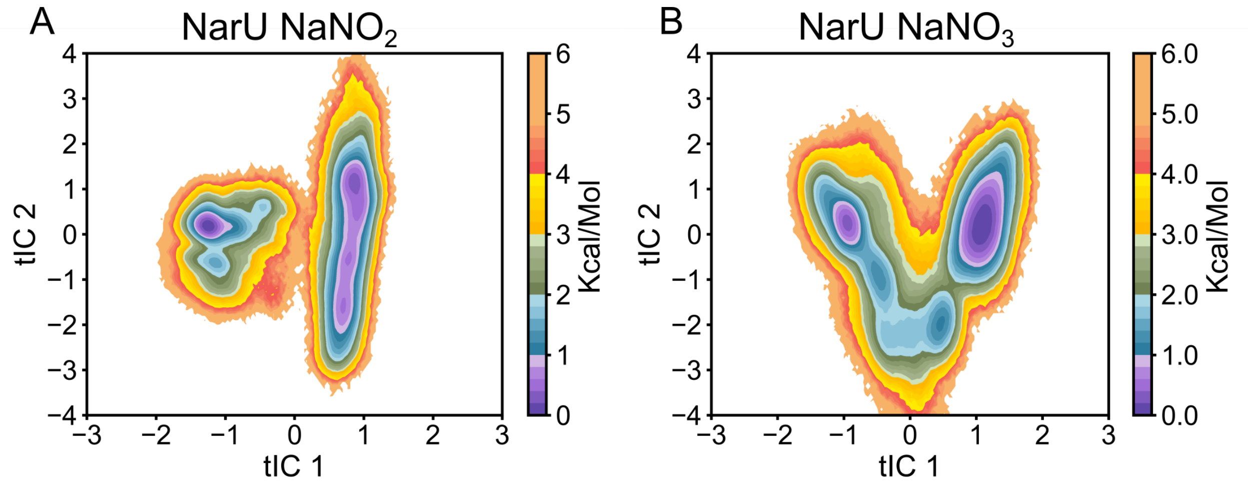


Figure S8: tICA plots projecting tIC 2 vs. tIC 1 for the A) $NO_2^-$ and B) $NO_3^-$ systems.

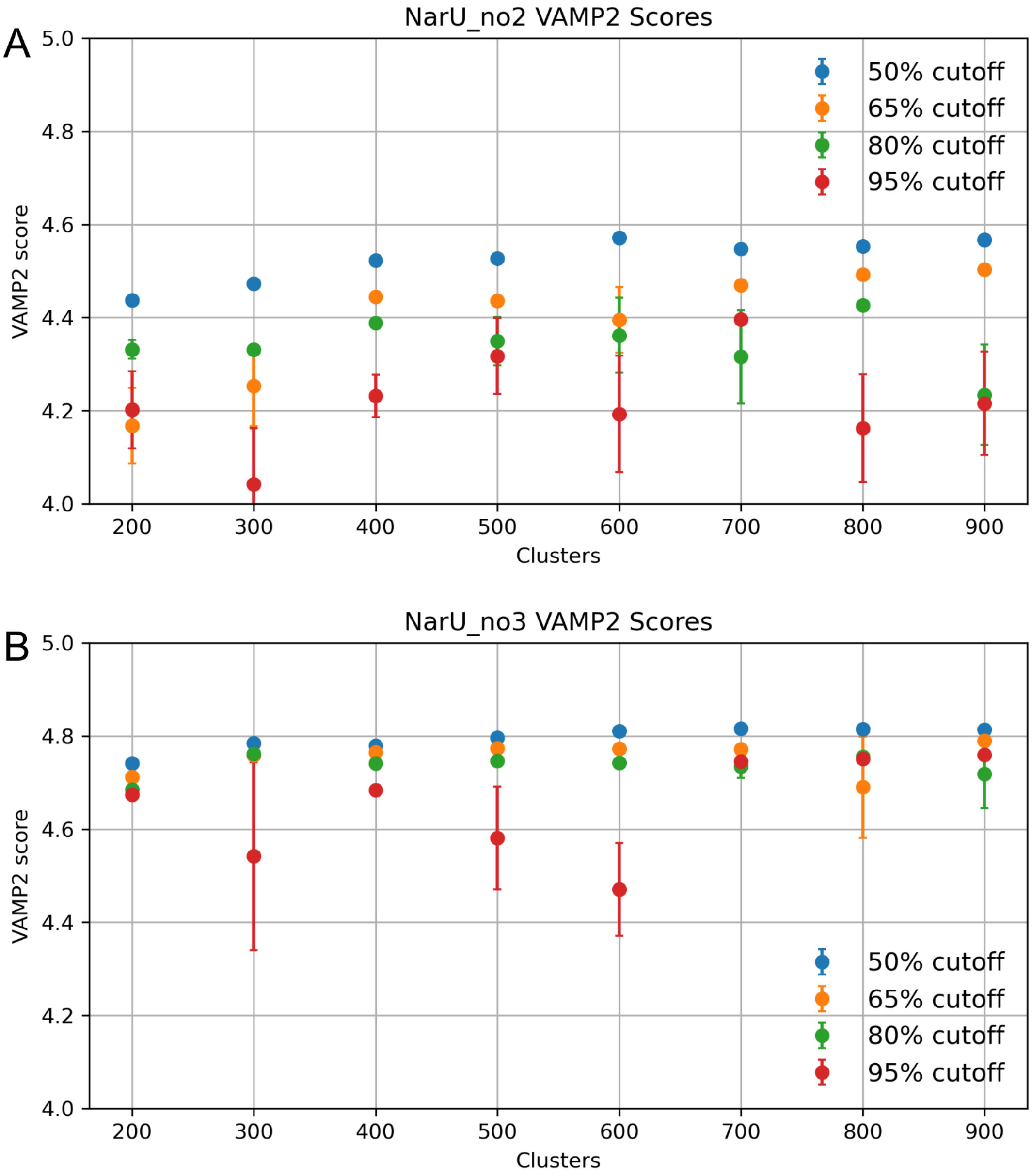


Figure S9: VAMP2 scores given differing cluster numbers and variance cutoffs for the A) $NO_2^-$ and B) $NO_3^-$ systems. Based on these scores, 900 clusters and 50% cutoff was used for $NO_2^-$ MSM construction while $NO_3^-$ used 800 clusters and 50% cutoff.

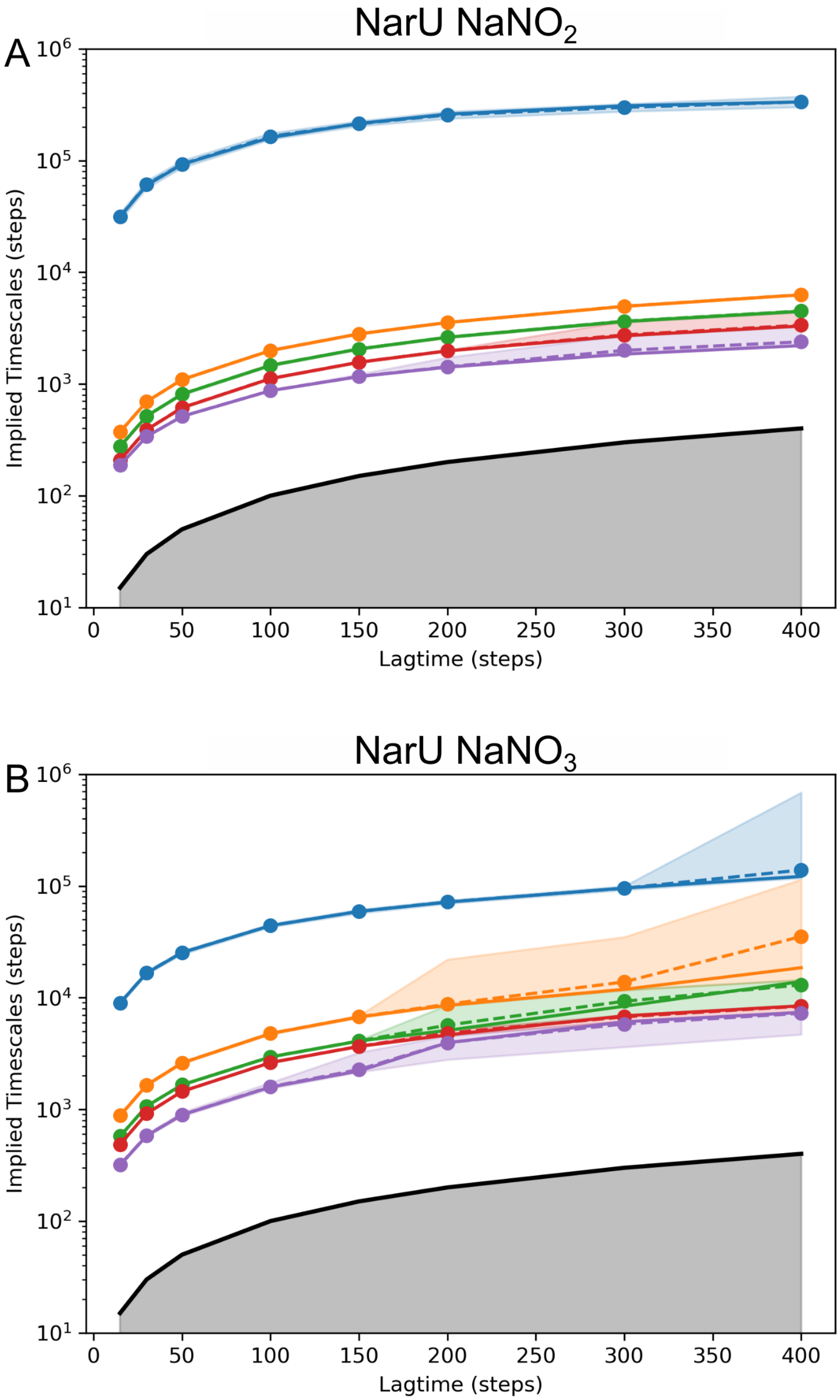


Figure S10: Implied timescale plots of the A) $NO_2^-$ and B) $NO_3^-$ systems. Based on these plots 150 steps, or 15 ns, was used as the lagtime for MSM construction in both systems.

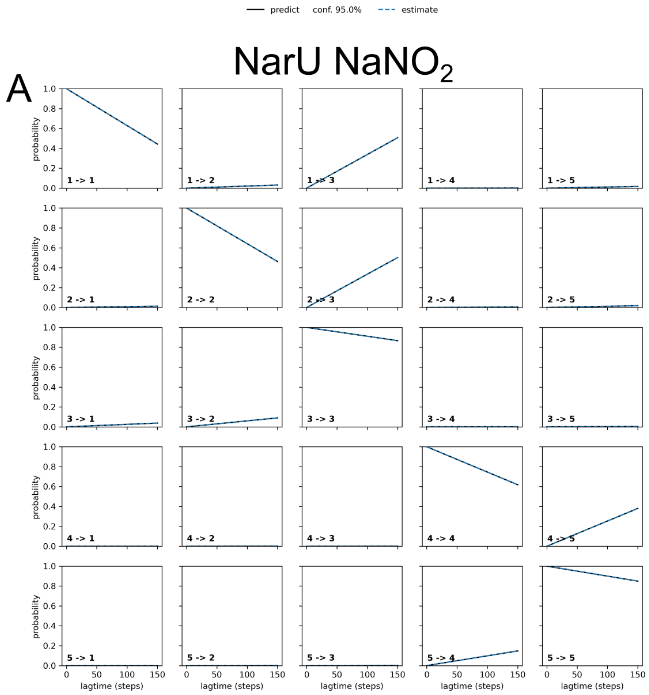


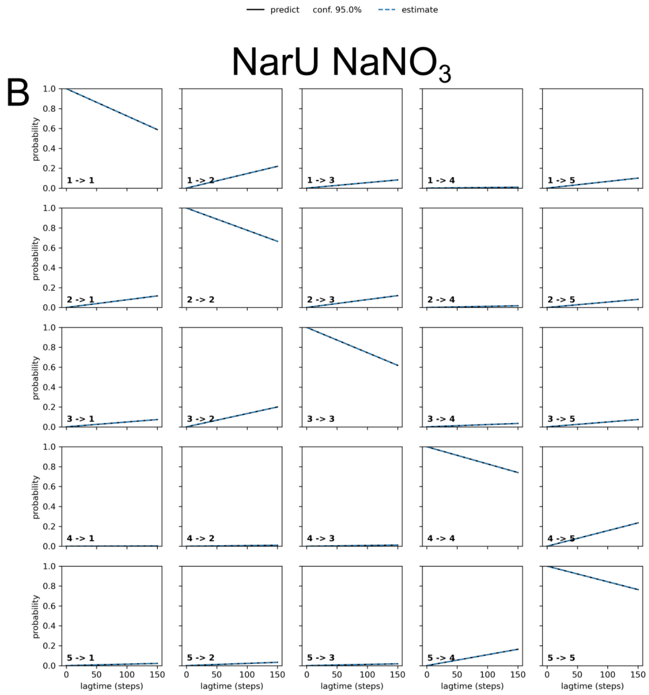


Figure S11: Chapman-Kolmogorov tests of a 5 macrostate system for the A) $NO_2^-$ and B) $NO_3^-$ systems.
